# Repeated Binge Alcohol Drinking Leads to Reductions in Corticostriatal Theta Coherence in Female but not Male Mice

**DOI:** 10.1101/2024.03.07.581791

**Authors:** Cherish E. Ardinger, Christopher C. Lapish, David N. Linsenbardt

## Abstract

Decreased functional connectivity between the striatum and frontal cortex is observed in individuals with alcohol use disorder (AUD), and predicts the probability of relapse in abstinent individuals with AUD. To further our understanding of how repeated alcohol (ethanol; EtOH) consumption impacts the corticostriatal circuit, extracellular electrophysiological recordings (local field potentials; LFPs) were gathered from the nucleus accumbens (NAc) and prefrontal cortex (PFC) of C57BL/6J mice voluntarily consuming EtOH or water using a ‘drinking-in-the-dark’ (DID) procedure. Following a three-day acclimation period wherein only water access was provided during DID, mice were given 15 consecutive days of access to EtOH. Each session consisted of a 30-minute baseline period where water was available and was followed immediately by a 2-hour period where sippers containing water were replaced with new sippers containing either unsweetened 20% (v/v) EtOH (days 4-18; DID) or water (days 1-3; acclimation). Our analyses focused primarily on theta coherence during bouts of drinking, as differences in this band are associated with several behavioral markers of AUD. Both sexes displayed decreases in theta coherence during the first day of binge EtOH consumption. However, only females displayed further decreases in theta coherence on the 14^th^ day of EtOH access. No differences in theta coherence were observed between the first and final bout on any EtOH drinking days. These results provide additional support for decreases in the functional coupling of corticostriatal circuits as a consequence of alcohol consumption and suggests that female mice are uniquely vulnerable to these effects following repeated EtOH drinking.

## Introduction

Previous work has reported that resting-state electroencephalograms (EEGs) of individuals with alcohol use disorder (AUD) show higher power in the theta band as compared to individuals without this diagnosis (Pollock et al., 1992, Mumtaz et al., 2017). Power is also higher in the theta band in individuals with a history of binge drinking as compared to non-binge drinkers (Lopez-Caneda et al., 2017, Affan et al., 2018). Preclinical studies in rodents also support the involvement of theta oscillations in excessive alcohol drinking (McCane et al., 2018, Henricks et al., 2019a, Henricks et al., 2019b). These findings suggest that theta oscillations in particular neural circuits may be critical intermediaries driving patterns of repeated alcohol consumption.

To date, numerous neural circuits have been implicated in alcohol consumption (Koob and Volkow, 2016, Koob and Volkow, 2010). However, the corticostriatal circuit in particular has been found to undergo functional adaptations that mediate a shift from occasional to higher-risk drinking (Belin et al., 2009, Barker and Taylor, 2014). Communication between corticostriatal brain regions, which can be measured as coherence, has been shown to be impaired following alcohol use (Courtney et al., 2013, Galandra et al., 2019). Recent evidence from a large multi-site collaborative study suggests coherence between brain regions in theta oscillations (i.e. theta coherence) may be key to understanding the relationship between alcohol drinking and neural function (Meyers et al., 2021). Theta coherence in the corticostriatal circuit is decreased in individuals diagnosed with alcohol use disorder (AUD), and this coherence is negatively correlated with time to relapse; i.e. less coherence was associated with a higher likelihood of a setback (Camchong et al., 2013). Further, in both rats and humans, adolescent alcohol exposure is associated with lower cortical theta coherence than control (Ehlers et al., 2020), suggesting that theta coherence within the corticostriatal circuit is important in alcohol use.

Previous research has also shown that neural activity in the prefrontal cortex (PFC) is crucial to encoding reward value (Hernandez and Moorman, 2020, Amarante et al., 2017, Amarante and Laubach, 2020). Theta coherence in the medial PFC was higher when rodents licked a high-value sucrose reward as compared to a low-value reward (Amarante et al., 2017, Amarante and Laubach, 2020). Increases in theta synchrony have been reported between the mPFC and nucleus accumbens (NAc) in alcohol-preferring (P) rats during alcohol drinking. However, less theta synchrony was observed in P rats over the total number of trials, presumably as a consequence of increased alcohol intake over the sessions (McCane et al., 2018). These results suggest increases in synchrony in the corticostriatal circuit during decision making (i.e. choosing to take a drink); however, the cumulative effect of multiple exposures to alcohol may cause these regions to become unsynchronized over time. Critical to the current paper, these findings suggest that theta synchrony in the corticostriatal circuit is influenced by both alcohol consumption (i.e. bouts) and alcohol drinking history.

Changes in corticostriatal theta power and coherence play a role in the development of AUD. However, there remains a need to systematically investigate the adaptations within corticostriatal theta neural activity which are induced by multiple alcohol drinking experiences to better understand how theta oscillations are impacted by sustained alcohol drinking. To begin to answer this question, the current study utilized extracellular electrophysiological recordings collected from the NAc and PFC of mice voluntarily consuming ethanol (EtOH) or water. Drinking bouts were time-locked with electrophysiological recordings, which allowed us to precisely evaluate neural oscillations as a function of drinking behavior. The primary goal of the current study was to identify how corticostriatal theta coherence, a measure of connectivity between brain regions, is influenced by repeated binge drinking. Thus, we evaluated changes in neural activity over many consecutive days of repeated EtOH intake, hypothesizing that theta coherence would progressively decrease.

## Methods

### Subjects

Data were gathered from 14 adult C57BL/6J female (*n*=7) and male (*n*=7) mice. However, two mice (1 F, 1 M) had off-target placements. These mice were removed from all analyses. This left us with a total of 6 mice per sex for data analysis, which are presented below. All mice were bred in the AAALAC-approved School of Science vivarium at Indiana University-Purdue University Indianapolis (IUPUI) using breeders supplied directly from Jackson Laboratories. Mice were group-housed with 3-4 same-sex siblings after weaning on postnatal day (PND) 21 in standard Allentown mouse cages (11 x 7 x 5-inch). One week prior to surgery, mice were transferred from the breeding colony to alternate vivarium locations. At the time of this transfer, mice were single-housed in slightly larger (12.5 x 7.5 x 7.5-inch) cages with taller/flat cage tops to prevent implant interference/impacts and were allowed to acclimate for one week prior to surgery on PND 84 (±12 days). Testing began following a one-week recovery period. Mice had *ad libitum* access to water and rodent chow (LabDiet 5001) at all times except during EtOH DID testing sessions when water was replaced with EtOH. Food remained available during DID. All procedures were approved by the IUPUI School of Science Animal Care and Use Committee and conformed to the Guidelines for the Care and Use of Mammals in Neuroscience and Behavioral Research (National Research Council, 2003).

### Surgery

Mice were implanted with custom-built electrodes made from 25 µm tungsten wire (California Fine Wires, Grover Beach, CA) inserted into a silica tubing matrix and then pinned to 16-channel Electrode Interface Boards (EIBs; Neuralynx, Bozeman, MT). Probes were configured such that 8 wires terminated in each of the two targeted brain regions - the Prefrontal Cortex (PFC: A/P: 1.94, M/L: ±0.5, D/V: -2.25) and Nucleus Accumbens (NAc: A/P: 1.10, M/L: ±0.8, D/V: -4.25). Anesthesia was induced with isoflourane at 3% in oxygen, and then maintained at 1-2% in oxygen during surgery. A subcutaneous injection of Ketoprofen (5 mg/kg, in a volume of 0.1 ml / 100g) was administered to reduce post-surgical pain. A sub-dermal injection of Bupivacaine (2.5 mg/mL, 0.05 mL) was administered directly over the incision site prior to exposing and cleaning the skull surface. Next, a unilateral craniotomy was made, electrodes were lowered and secured with dental acrylic, and two ground wires attached to the EIB and affixed with stainless steel screws were placed over the cerebellum. A final head cap was then completed using the same dental acrylic to ensure all components were covered and any burrs were removed. Mice were monitored closely for one-week following surgery, including daily checks for weight, feeding, and signs of distress. Cephazolin (30mg/kg SC, 3 mg/mL) and/or a topical antibiotic (2% bacitracin/polymixin) was administered as necessary during this recovery period. During the first 2-3 days of recovery, mice were given LabDiet 5015 mixed with water as a high protein ‘treat’ to encourage post-operative feeding. This treat was available in addition to their standard ad libitum diet, LabDiet 5001. After this one-week recovery period, all mice were at or above pre-surgery weight and had fully recovered. Local field potentials (LFPs) were then recorded during DID testing (see below).

### Solutions

EtOH for drinking experiments was prepared by diluting 190 proof EtOH from Pharmco, Inc. (Brookfield, CT) to 20% v/v in tap water. Water for water drinking days was obtained from the same tap as water that was used to dilute the EtOH. Drinking solution was prepared at the beginning of the experiment and stored in a sealed container. This prepared solution was used to fill ball-bearing sipper tubes during DID procedures described below.

### Behavioral Electrophysiology Recording System

Home cages were located directly beneath overhead frictionless motorized rotary joints that allowed tethered subjects unrestricted movement during recording sessions. Rotary joints were connected to an OpenEphys recording system, which was in turn connected to a data acquisition computer. The acoustics of ball-bearing sipper tubes during drinking bouts were detected using piezo microphones, amplified via a professional audio soundboard, and relayed to an I/O board integrated with the OpenEphys acquisition systems tertiary analog inputs via an HDMI cable. This configuration allowed for precise alignment of bouts to neurophysiology data. The X and Y Cartesian coordinates of mice within the home cage were recorded with AnyMaze tracking software, were then converted to voltage using an AnyMaze digital-to-analog converter, and similarly time-locked with neurophysiology data as drinking bouts.

### Behavioral Electrophysiology Recording Procedures

Mice underwent a common home-cage binge drinking procedure (drinking-in-the-dark, DID) (Rhodes et al., 2005) in which standard home cage water bottles are replaced with a single custom-built double ball-bearing sipper tube containing either 20% EtOH or water three hours into the dark cycle. The first three days mice were given two hours of access to water in these tubes. The following 15 days mice were given access to EtOH. The final 3 days were identical to the first 3 days (water access). Thus, 21 total days of testing were conducted for each subject (see *Figure 1*).

**Figure 1.**
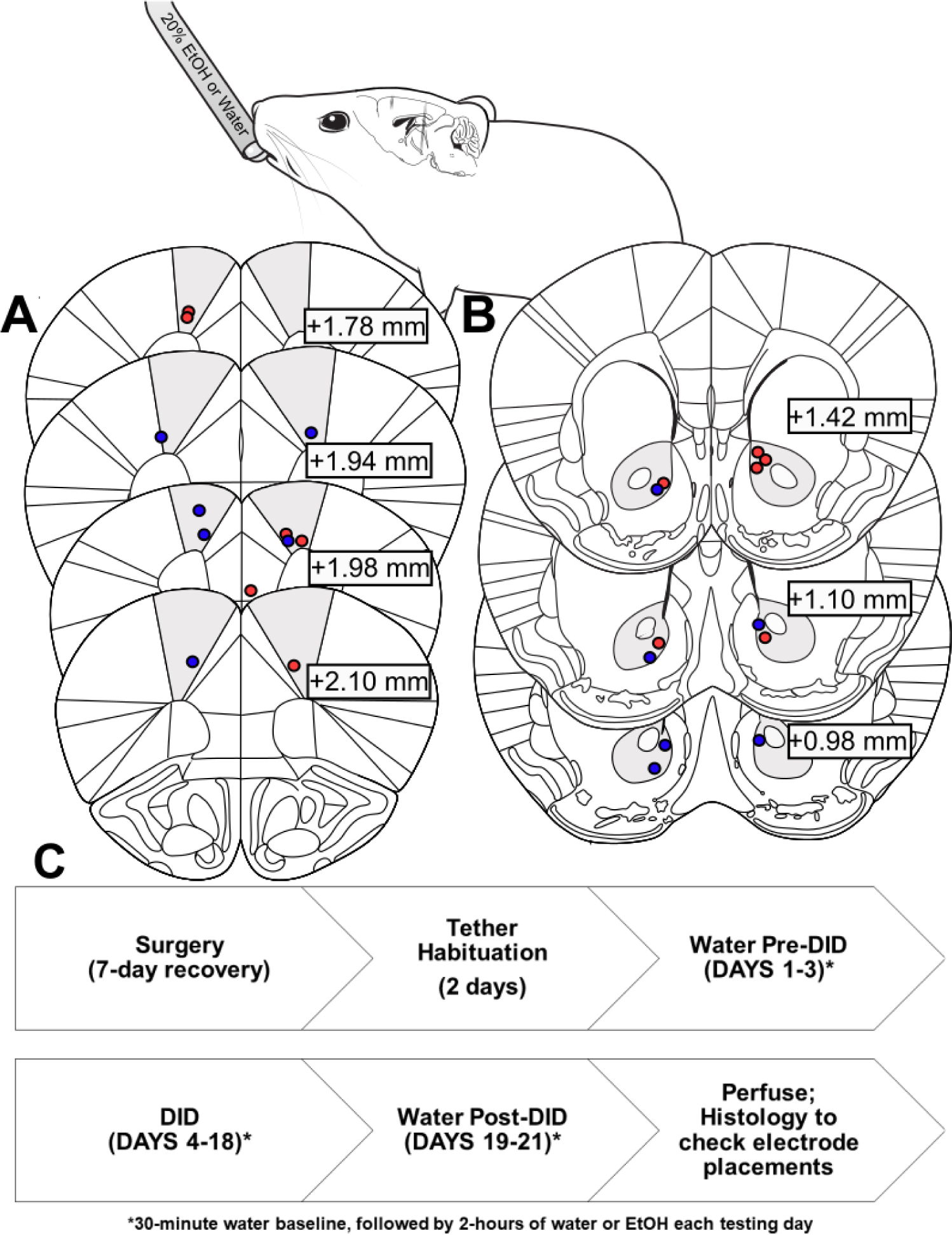
Overview of experiment and placements. Placements for females (red) and males (blue) are shown in the PFC (A) and NAc (B). A timeline of experiments is shown in (C). The mouse drinking cartoon was modified from scidraw.io (Tyler & Kravitz, 2020). Placements were determined using Paxinos and Franklin’s Mouse Brain in Stereotaxic Coordinates 3^rd^ edition Atlas.

On each testing day, one hour into the dark cycle, mice were transferred from the colony room to the electrophysiology testing room and given 1.5 hours to acclimate in the location where electrophysiology recordings were subsequently collected. Mice were then tethered to a rotary joint and data acquisition was initiated for 30-minute baseline recordings. During baseline recordings, home-cage water bottles were replaced with standard ball-bearing sippers, which always contained water. The end of this 30-minute baseline period was immediately followed by a 2-hour DID testing session. Water-containing baseline ball-bearing sippers were replaced with new identical sippers containing EtOH (days 4-18) or water (water pre-DID days 1-3; or water post*-*DID days 19-21). The first two days served as ‘tether habituation’ days; data from tether habituation is not shown, and these days were not included in the three water pre-DID days. Fluid intake was recorded from volume left in the sippers at the end of baseline (water only) and DID session (water or EtOH, depending on the testing day).

### Immunohistochemistry

When testing was completed, mice were anesthetized with isoflurane as described in the surgery section above and a stimulus isolator was used to create lesions at the site of each of the wires via sending a 1 mA current through the EIB into each wire for one second. Twenty-four hours later mice were transcardially perfused and brains were removed to verify electrode placements. Mice were irreversibly anesthetized with urethane via intraperitoneal injection at a dose of 1.5 g/kg dissolved in sterile water in a volume of 1.0 mL/kg, and transcardially perfused with isotonic saline followed by 4% paraformaldehyde (PFA). Brains were removed and placed in PFA for 4 days and then transferred to sucrose solution (30%) for cryopreservation. Standard slicing and staining procedures using DAPI (Santa Cruz, Cat No: sc-3598) and GFAP (Fisher, Cat No: NBP105198) were followed to determine electrode placement. Placements can be seen in Figure 1.

### Behavioral Analyses

Locomotor activity was analyzed using a 2 (sex) x 21 (days) mixed methods 2-way analysis of variance (ANOVA). Water intake and EtOH intake were analyzed separately using 2 (sex) x 3 (water pre/post DID days) or 15 (EtOH DID days) mixed methods 2-way ANOVAs in GraphPad Prism. Animals were removed from analyses on a given day if they had a sipper leak or had faulty video / locomotor tracking. These technical issues were monitored closely and were generally able to be resolved the following day, but explain the variance in degrees of freedom between locomotor and intake analyses.

### Electrophysiology Pre-Processing

Data were imported into MATLAB using open-source MATLAB functions (available on the <u>OpenEphys GitHub page</u>), down sampled to 1000Hz, and normalized via median subtraction to minimize potential impacts of motion artifacts and volume conduction. Prior to preprocessing, excessively noisy channels were identified as those with voltages greater than two standard deviations above the mean of the other channels within a given brain region and removed.

### Power and Coherence Analyses

The primary focus of the current analyses was to assess neural activity associated with alcohol consumption (i.e. drinking bouts), and how this consumption-related activity changed following many alcohol drinking experiences. Bouts were defined identical to Darevsky et a. (2019): three or more licks that occur ≤ 1 sec apart. Bouts were manually identified and curated by reviewing microphone data streams and high-resolution video recordings. LFP data ± 4 seconds around the onset of each bout were extracted from each brain region separately, averaged, and assessed using the MATLAB ‘spectrogram’ function (500 ms window; 450 sample overlap; 1000 Hz sample rate). Coherence of the same bout-aligned LFP epochs detailed above was assessed using the MATLAB function ‘wcoherence’.

Given our *a priori* hypotheses of theta band activity, data surrounding bouts on days three (Water baseline), four (D1 EtOH), and seventeen (D14 EtOH) within this band was averaged and evaluated using 3-way (sex x brain region x day; for theta power) or 2-way (sex x day; for theta coherence) ANOVAs. Results revealed significant interactions with sex, described below. To best understand the within sex effects, separate one-way ANOVAs followed by Tukey’s multiple comparison tests were then conducted. Day 17 (D14 EtOH) was selected instead of Day 18 (D15 EtOH, the last day of EtOH drinking) because 40 uL of retro-orbital blood was drawn (or attempted to be drawn) for blood ethanol concentration (BEC) analyses immediately following the end of the DID session on day 17. The head-cap required for the chronic awake-behaving recordings made this blood draw procedure more challenging than is typical. We were unable to obtain blood samples from some of the mice, which is why BEC data are not presented. To ensure that any association of drinking with the experience of this procedure did not impact results, day 17 (day 14 EtOH) was chosen for analyses.

However, we recognize that BEC data are important in preclinical alcohol studies. Therefore, we have estimated BECs using a regression equation from prior data in our laboratory from C57Bl/6J mice undergoing a similar procedure (14 days of 2-hour DID) (Linsenbardt and Boehm, 2014). These results are reported below.

## Results

### Intake and Locomotion over days

Analysis of 2-hour water intake across the first 3 days of testing (water pre-DID) identified no main effect of day [*F* (1.826, 15.52) = 1.24, *p* > 0.05], sex [ *F* (1, 17) = 1.91, *p* > 0.05], or interaction of day by sex [*F* (2, 17) = 0.15, *p* > 0.05], (Figure 2A). Similarly, intake during the EtOH DID days revealed no main effect of day [*F* (4.401, 32.69) = 2.35, *p* > 0.05], sex [*F* (1, 10) = 0.15, *p* > 0.05], or interaction of day by sex [*F* (14, 104) = 0.64, *p* > 0.05], (Figure 2A). Analysis of 2-hour water intake across the last 3 days of testing (water post-DID) also identified no main effect of day [*F* (2, 19) = 0.61, *p* > 0.05], sex [*F* (1, 19) = 0.06, *p* > 0.05], or interaction of day by sex [*F* (2, 19) = 0.22, *p* > 0.05], (Figure 2A). Lastly, analysis of locomotor activity identified no main effect of day [*F* (3.41, 25.89) = 1.01, *p* > 0.05], sex [*F* (1, 10) = 2.15, *p* > 0.05], or interaction of day by sex [*F* (20, 152) = 0.49, *p* > 0.05], (Figure 2B).

**Figure 2.**
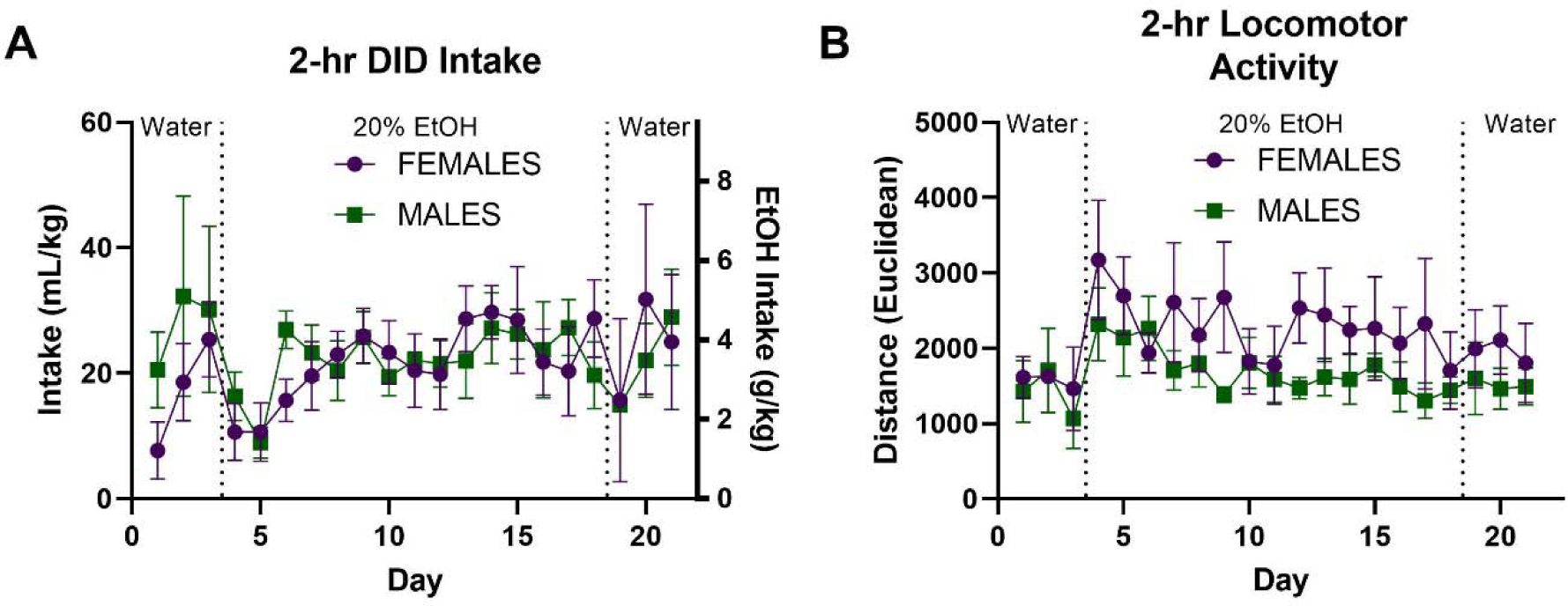
Average intake (A) and locomotor activity (B). There were no main effects of sex, day, or interaction of day by sex when analyzing intake or locomotion.

### Estimated BEC

A regression equation generated from the relationship between intake and BEC in 2-hour DID from C57BL/6J mice who underwent DID for 14 days using similar methodology as the current study (Linsenbardt and Boehm, 2014) was used to estimate BECs for mice in the current study. The equation, Y = 48.75*Intake(mL/kg) - 26.79, estimated average BECs of our mice to be 133.07 (± 38.41) and 171.27 (± 24.41) mg/dL on day 14 for females and males, respectively. Data are presented as mean ± SEM.

### Theta Power

Average drinking-bout-associated theta power during water consumption, the first alcohol consumption session, and the 14^th^ alcohol consumption session can be seen in Figure 3. A 3-way ANOVA revealed significant main effects of sex [*F* (1, 834) = 197.3, *p* < 0.0001], day [*F* (2, 834) = 6.21, *p* < 0.001], and brain region [*F* (1, 834) = 42.48, *p* < 0.0001], as well as significant day x sex [*F* (2, 834) = 14.45, *p* < 0.0001], day x brain region [*F* (2, 834) = 16.83, *p* < 0.0001], and sex x brain region [*F* (1, 834) = 12.27, *p* < 0.001] interactions. To better understand what was driving these interactions, we then conducted separate one-way ANOVAs with each brain region and sex (see below).

**Figure 3.**
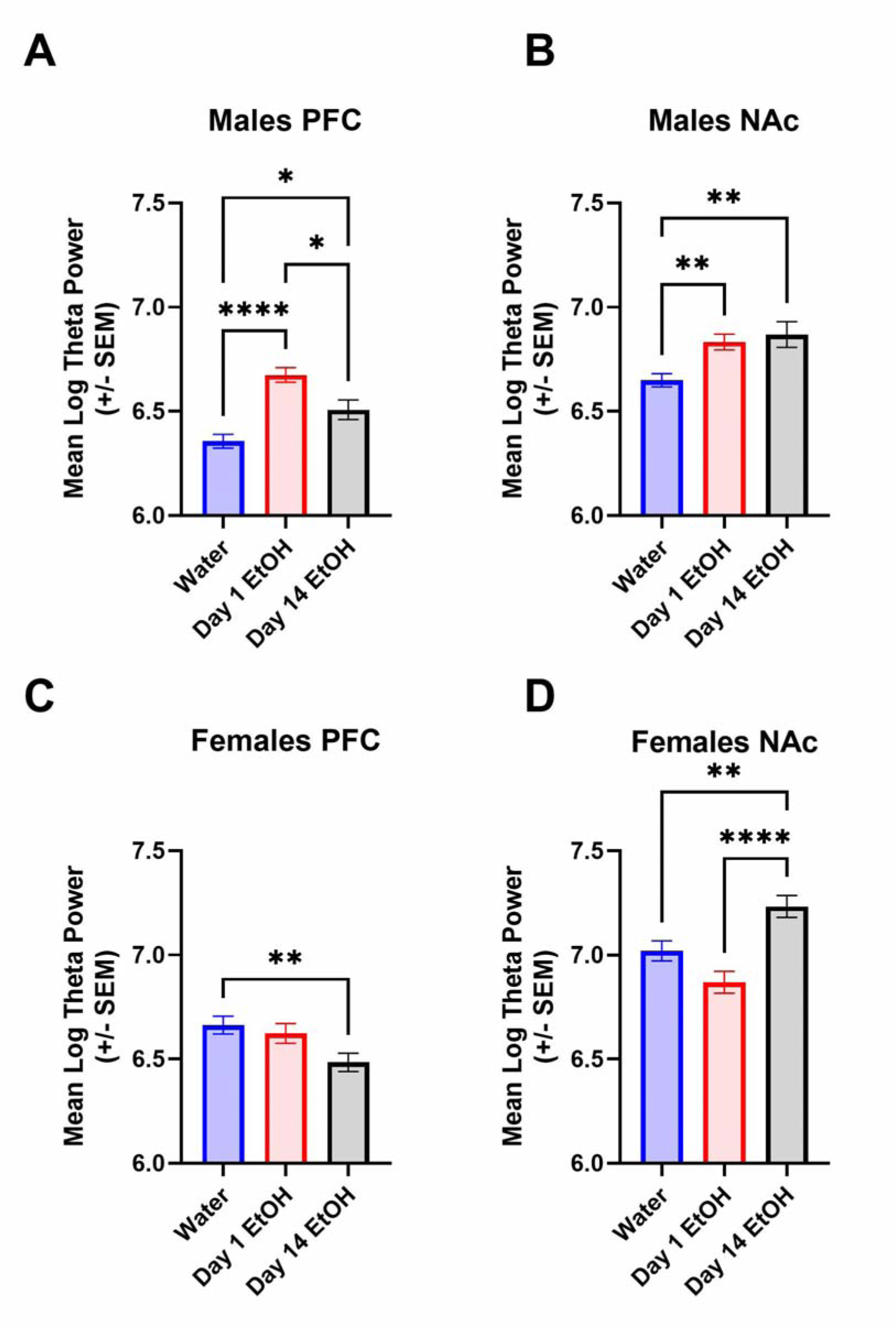
Theta Power During Binge Drinking Differs Between Females and Males. In males, in both the PFC (A) and NAc (B) during drinking bouts, theta power is higher when mice consumed EtOH as compared to water. In the PFC, the first binge drinking day produced higher theta power as compared to both the water and 14^th^ day of DID. In females, in the PFC, there were no differences between days (C). In the NAc, theta power was significantly higher on day 14 of DID as compared to both water and day 1 of DID (D).

In the male PFC (Figure 3A), alcohol drinking significantly increased theta power [*F* (2, 237) = 22.47, *p* < 0.0001], with Tukey’s post-hoc tests confirming significantly higher theta power on the first (*p* < 0.0001) and 14^th^ alcohol days (*p* < 0.05) versus the water session, as well as higher theta power on the first alcohol day compared to the 14th (*p* < 0.05). A similar pattern was observed in males in the NAc [*F* (2, 237) = 8.74, *p* < 0.0001], with Tukey’s post-hoc tests indicating significantly lower theta power during bouts in the water session versus the first (*p* < 0.001) and 14^th^ alcohol sessions (*p* < 0.001; Figure 3B). Thus, in males, theta power was higher in the PFC and NAc during the first alcohol drinking session, but this effect decreased following 2 weeks of alcohol access in the PFC, whereas it remained increased in the NAc.

In the female PFC (Figure 3C), alcohol drinking significantly decreased theta power [*F* (2, 180) = 4.79, *p* < 0.01], with Tukey’s post-hoc tests confirming significantly lower theta power on the 14th alcohol day versus the water session (*p <* 0.001). Conversely, in the NAc of females, 14 days of alcohol experience significantly increased theta power [*F* (2, 180) = 11.76, *p* < 0.0001], and Tukey’s tests confirmed this increase was significant between both the water session (*p* <0.01) and the first alcohol session (*p* < 0.0001) (Figure 3D).

### A Single Alcohol Drinking Experience Decreases Theta Coherence in Both Sexes, and is Further Decreased Following Repeated Experiences in Females Only

Average drinking-bout-associated theta coherence during water consumption, the first alcohol consumption session, and the 14th alcohol consumption session can be seen in Figure 4. First, a two-way sex (2: female vs. male) x day (3: water vs. day 1 EtOH vs. day 14 EtOH) ANOVA revealed a significant interactions of sex x day [*F* (2, 417) = 5.48, *p* < 0.01] and main effects of sex [*F* (1, 417) = 32.51, *p* < 0.0001], and day [*F* (2, 417) = 17.68, *p* < 0.0001]. To better understand what was driving these interactions, we then conducted separate one-way ANOVAs with each brain region and sex (see below).

**Figure 4.**
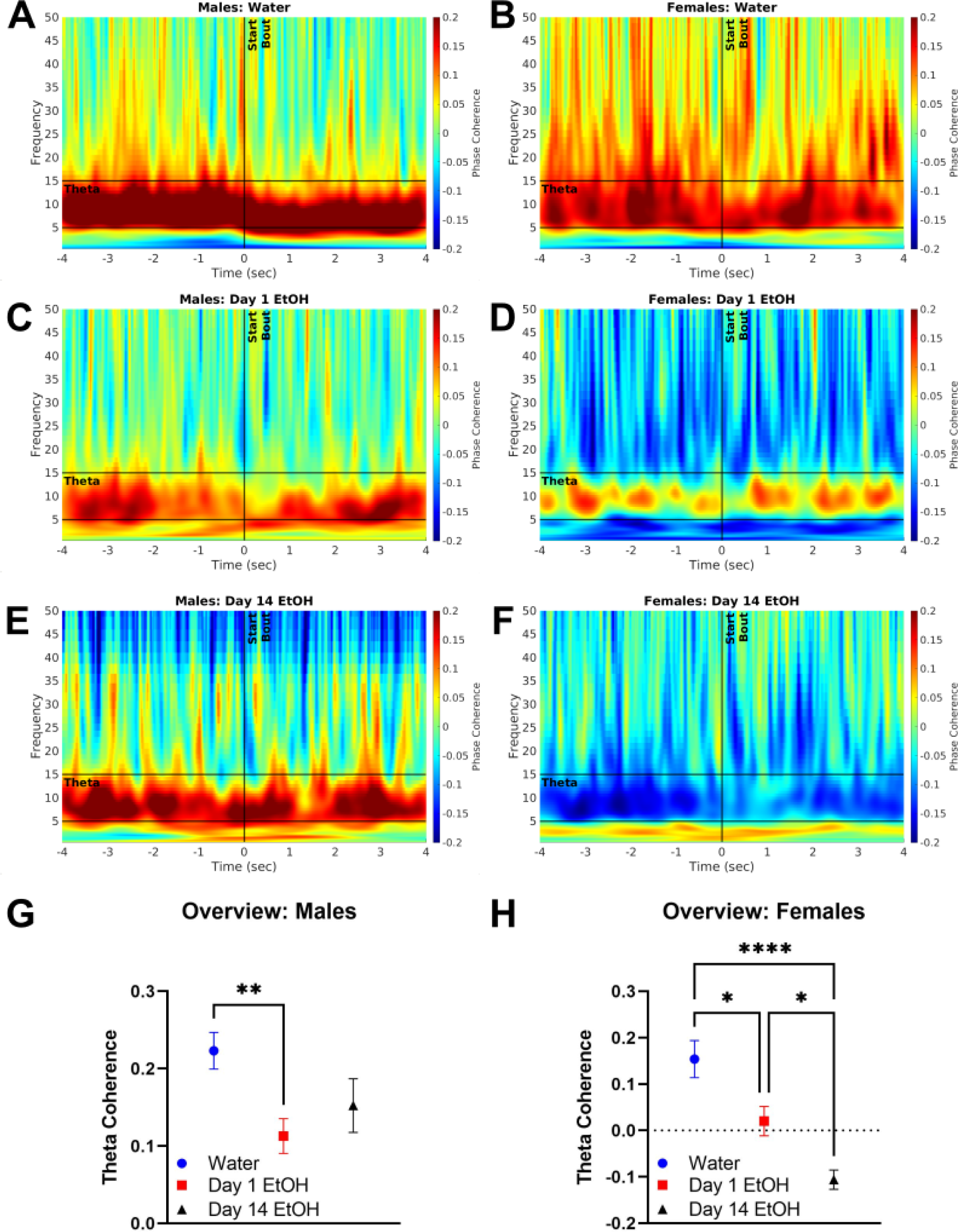
Repeated Binge Drinking Decreases Theta Coherence in Females. In males, there was a decrease in theta phase coherence on the first day of DID (C) as compared to water (A). This decrease was not observed on day 14 of DID (E); summary (G). In females, there was a binge drinking experience-dependent decrease in theta phase coherence, where significant differences were observed between water (B) and days 1 (D) and 14 (F), and days 1 and 14; summary (H).

In males, significant alterations in theta phase coherence during bouts [*F* (2, 237) = 5.57; *p* < 0.01] were confirmed by post-hoc tests as being driven by significant decreases during the first alcohol session (*p* < 0.01), which recovered to the levels of the water session by the 14^th^ day of DID (Figure 4, left panels).

In females, significant alterations in theta phase coherence during bouts [*F* (2, 180) = 17.20; *p* < 0.0001] were confirmed by post-hoc tests as being driven by significant decreases during the first alcohol session (*p* < 0.05) and more prominent decreases in the 14^th^ alcohol sessions (p<0.0001) (Figure 4, right panels).

Importantly, this decreased theta coherence was specific to alcohol drinking. There were no significant differences in coherence between the first alcohol day and the 14^th^ alcohol day during bouts which occurred in the 30-minute water baseline period. This was true for both males, *t* (82) =0.13, *p* > 0.05, and females, *t* (36) =0.87, *p* > 0.05 (null data not shown). Lastly, to assess how EtOH’s pharmacological actions over a given 2-hour session influenced coherence, coherence was compared from the first bout to the last bout. Separate t-tests were conducted for females and males on day 1 and day 14 of EtOH and found no differences between the first and last bout on either day for either sex.

There is an established relationship in the literature between theta oscillations and locomotor activity in rodents (Buzsáki et al., 1983, Chen et al., 2011, Long et al., 2014, Kropff et al., 2021). To assess if our observations were influenced by activity required to approach the sippers, we compared locomotion time-locked to theta coherence (Figure 5). We found no evidence of a relationship between these 2 measures on any of the 3 days of focus; theta coherence remained stable regardless of fluctuations in locomotion.

**Figure 5.**
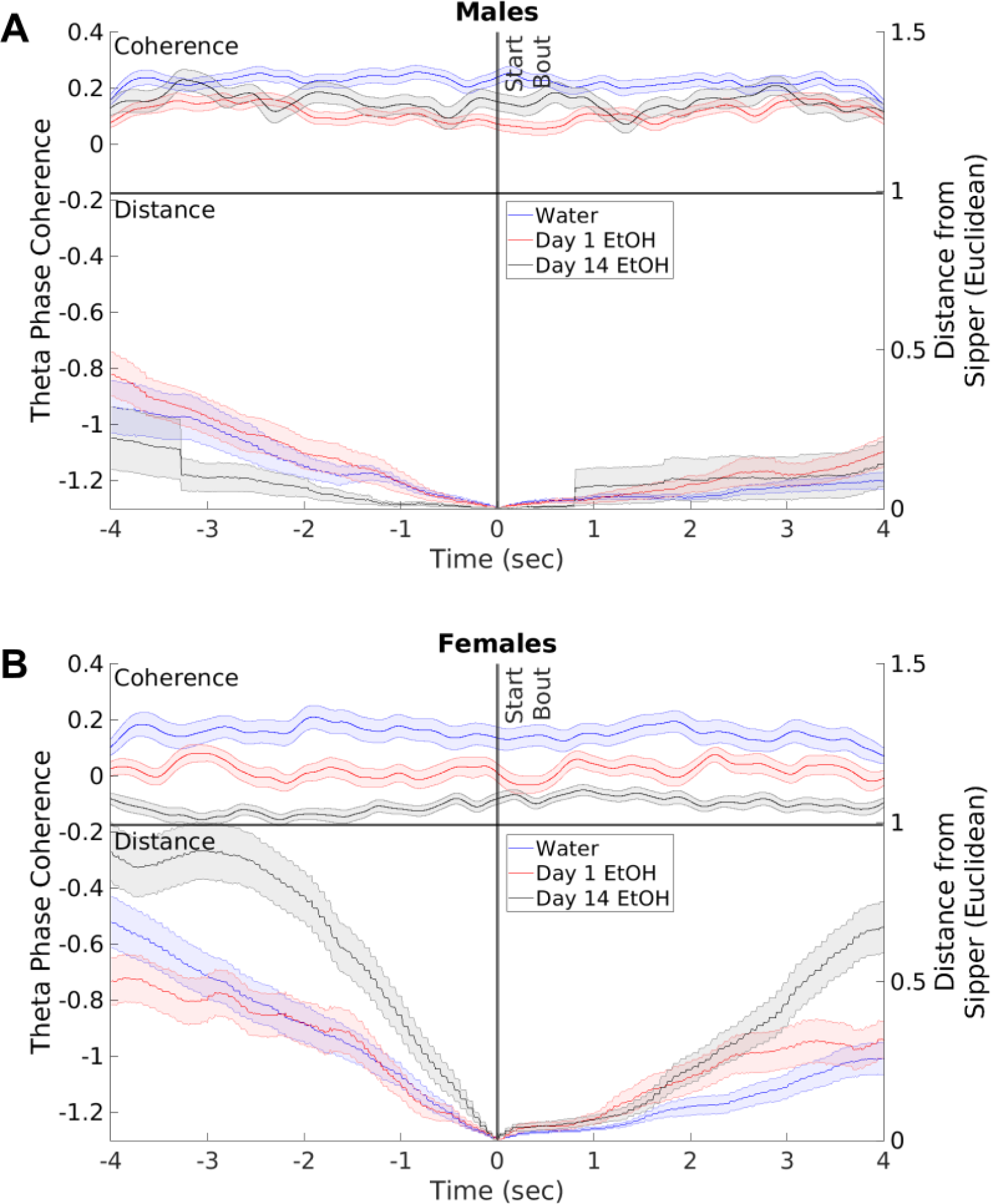
Coherence is Not Likely Influenced by Sipper Approach. Theta coherence near bout start times does not appear to be influenced by sipper approaches and retreats in males (A) or females (B).

## Discussion

The results of this work reveal sex-dependent differences in theta oscillations associated with alcohol drinking. Males displayed an alcohol-induced increase in corticostriatal theta power on the first day of EtOH (Figures 3A, 3B), whereas no differences in theta power were observed in females on this day (Figures 3C, 3D), indicating that corticostriatal circuits are more sensitive to EtOH in EtOH-naïve males vs EtOH-naïve females. By the 14^th^ day of EtOH consumption, both sexes displayed significant differences in theta power versus water, with males displaying persistent increases relative to water sessions (Figures 3A, 3B) and females displaying bidirectional alterations; decreases in theta power in the PFC and increases in theta power in the NAc (Figures 3C, 3D). The most surprising observation however was significantly lower theta coherence on the 14th EtOH access day in females but not males (Figure 4H). We confirmed that this was EtOH-drinking specific and not due to generally lower theta coherence over days through assessment of the 30-minute water baseline recordings prior to DID EtOH access. Collectively, these results are consistent with findings of higher theta power in individuals with a history of alcohol drinking (Lopez-Caneda et al., 2017, Pollock et al., 1992, Rangaswamy et al., 2003, Affan et al., 2018), but emphasize important differences that may exist as a function of brain region and sex.

An important limitation of the current study is the lack of a sucrose / saccharin control group. It is possible that, alternatively or additionally, the increased theta power in males, and decreased theta power in males and females, on day 1 may be driven by the novelty of the alcohol. A relationship between novelty seeking and AUD is well-described (Manzo et al., 2014, Flagel et al., 2014; for a review, see Wingo et al., 2016). Studies have shown that licking sucrose tends to produce increases in both theta power (Wingerden et al., 2010) and theta coherence (Amarante et al., 2017, Amarante and Laubach, 2020, Horst and Laubach, 2013), suggesting that our day 1 coherence results may be specific to alcohol. Future work along the lines of the current study should include a sucrose / saccharin control to assess the impact of novelty on theta oscillations.

The interpretation of the sex differences detailed above are somewhat difficult to fit within the context of the existing literature, as most awake-behaving electrophysiology studies of voluntary alcohol consumption in rodents have only included male subjects (Morningstar et al., 2020, Henricks et al., 2019a, Hernandez and Moorman, 2020, Timme et al., 2022, McCane et al., 2018, Linsenbardt and Lapish, 2015, Linsenbardt et al., 2019). However, a recent study found that an acute 1 g/kg alcohol exposure decreased gamma power in the basolateral amygdala in females, whereas the same treatment decreased beta power in males (DiLeo et al., 2022). Another recent study in rats found that PFC and NAc oscillations could successfully predict intake in males, but not females (Henricks et al., 2019b), and that this prediction was influenced by estrous phase (Henricks et al., 2019b). Although estrous cycle was not tracked in the current work, future studies investigating the role of hormonal fluctuations to alcohol consumption and neural oscillations are warranted.

The rodent PFC and NAc are heterogenous structures that each contain subdivisions with differences in anatomy (e.g. afferent/effect projections) and biochemistry (e.g. neurotransmitters, receptors). In the current study, electrodes were primarily in the M2 region of the PFC, which send projections to the NAc (Li et al., 2018). To our knowledge, this is the first assessment of this projection in a rodent model of AUD. Of particular importance to relapse models is the projection from the dorsomedial PFC (prelimbic and anterior cingulate cortex) to the NAc core (Stefanik et al., 2013). Plasticity in this pathway plays a key role in driving reinstatement of drug seeking, when cued by a conditioned stimulus (McGlinchey et al., 2016). Furthermore, reductions in cue-evoked oscillations in the dorsomedial PFC are associated with reductions in alcohol seeking and drinking (McCane et al., 2018). However, these studies have been largely limited to male rodents. Therefore, it will be important to determine if the sex differences in theta coherence observed herein are observed in the dorsomedial PFC and NAc core.

An important next step in this line of research will be to determine the potential causality of our observations, for example, by evaluating the impact of experimental manipulations of theta coherence on alcohol consumption. To this end, a recent study in individuals with AUDs demonstrated that transcranial magnetic theta burst stimulation in the medial PFC led to a 3-fold increase in the probability of maintaining sobriety than those who received sham treatment (McCalley et al., 2022). Thus, we are currently well positioned to use the model we describe here to validate promising clinical treatments as well as to identify novel intervention strategies.

In summary, the current findings provide further support for decreased corticostriatal theta synchrony and increased corticostriatal theta power as biomarkers of alcohol drinking and/or AUD.

